# Predictive Systems Biology Modeling: Unraveling Host Metabolic Disruptions and Potential Drug Targets in Acute Viral Infections

**DOI:** 10.1101/2023.07.24.550423

**Authors:** Gong-Hua Li, Feifei Han, Rong-Hua Luo, Peng Li, Chia-Jung Chang, Weihong Xu, Xin-Yan Long, Jing-Fei Huang, Yong-Tang Zheng, Qing-Peng Kong, Wenzhong Xiao

## Abstract

**Background:** Host response is critical to the onset, progression, and outcome of viral infections. Since viruses hijack the host cellular metabolism for their replications, we hypothesized that restoring host cell metabolism can efficiently reduce viral production.

**Results:** Here, we present a viral-host Metabolic Modeling (vhMM) method to systematically evaluate the disturbances in host metabolism in viral infection and computationally identify targets for modulation by integrating genome-wide precision metabolic modeling and cheminformatics. We applied vhMM to SARS-CoV-2 infections and identified consistent changes in host metabolism and gene and endogenous metabolite targets between the original SARS-COV-2 and different variants (Alpha, Delta, and Omicron). Among six compounds predicted for repurposing, *methotrexate, cinnamaldehyde*, and *deferiprone* were tested *in vitro* and effective in inhibiting viral production with IC50 less than 4uM. Further, an analysis of real-world patient data showed that cinnamon usage significantly reduced the SARS-CoV-2 infection rate with an odds ratio of 0.65 [95%CI: 0.55∼0.75].

**Conclusions:** These results demonstrated that vhMM is an efficient method for predicting targets and drugs for viral infections.

## Background

Viral infections have consistently been a significant health concern throughout human society [1, 2]. Despite intensive research, a few drugs are FDA-approved to date for viral diseases[3-5]. The emergence of viral variants has also significantly impacted the efficacy of available therapeutics and vaccines[6-9]. There is still a critical need for developing new agents to treat viral infectious diseases. While current small-molecule drugs target viral proteins, investigations into the host pathways provide opportunities for discovering new therapeutic agents with different mechanisms that are either efficacious alone or in combination therapy. Since viruses hijack host cellular metabolism for replications, which require high energy expenditure and a large number of host metabolites[10] for the production of viral components, such as proteins, RNA, and lipids, we hypothesize that restoring host cell metabolism can efficiently negate viral production.

Genome-scale metabolic modeling can capture metabolic states of cells or tissues[11] and has been applied to understanding the mechanisms of diseases, such as obesity[12], NAFLD [13], cancer[14], and Alzheimer’s disease[15]. It has also been utilized to study human cell metabolic changes after viral infection and infer gene targets[16-18]. However, how to directly predict candidate drugs for viral infections by targeting the host metabolic response remains challenging.

We recently developed an algorithm of precision metabolic modeling (GPMM) by quantitative integration of the enzyme kinetics information from knowledge bases and enzyme levels from transcriptome and proteome data[19]. Together with the *in silico* genome-wide gene and metabolite knock-in/knock-out capability[20], this approach allows systematic evaluations of genes and endogenous metabolites as candidate targets to modulate the host metabolism. In addition, recent developments in chemoinformatics have enabled computational drug discovery by comprehensive comparison of molecular structures[21, 22] and integrated bioactivity data between a large number of small molecule compounds and the candidate targets[23].

Here we integrate these algorithms to unravel the host metabolic disruptions in viral infection and discover potential antiviral targets. We applied this method to analyze the genomic data of human cells infected by SARS-CoV-2 and identified gene and metabolite targets as well as drug candidates that inhibit both the original virus and its variants. We performed *in vitro* validation experiments on three out of six predicted drug candidates and found that they can effectively inhibit viral production in cell lines. Further, an analysis of real-world data from five major hospitals showed that cinnamon usage significantly reduced the risk of COVID-19 infection. Taken together, this computational approach is effective in aiding the modulation of host metabolism against viral infection.

## Results

### viral-host Metabolic Modeling (vhMM) for modulating host metabolism against viral infection

vhMM was developed to predict drug candidates to modulate host metabolism against viral infection by integrating several procedures, including system-level precision metabolic modeling, *in silico* system-wide knock-out analysis of genes and metabolites on metabolic fluxes, and chemoinformatics to identify drugs targeting these genes and metabolites for repurposing.

As shown in **Figure 1**, a metabolism network model of human cells after viral infection was first constructed that integrated human reconstructed metabolic models (e.g. Recon3D) and the COVID-19 disease map[24]. In total 28 SARS-CoV-2-specific proteins, negative and positive RNAs, and lipids were included in the curated virus-host metabolism map, along with the viral biomass reaction using protein and RNA compositions from the literature[24, 25].

**Figure 1:**
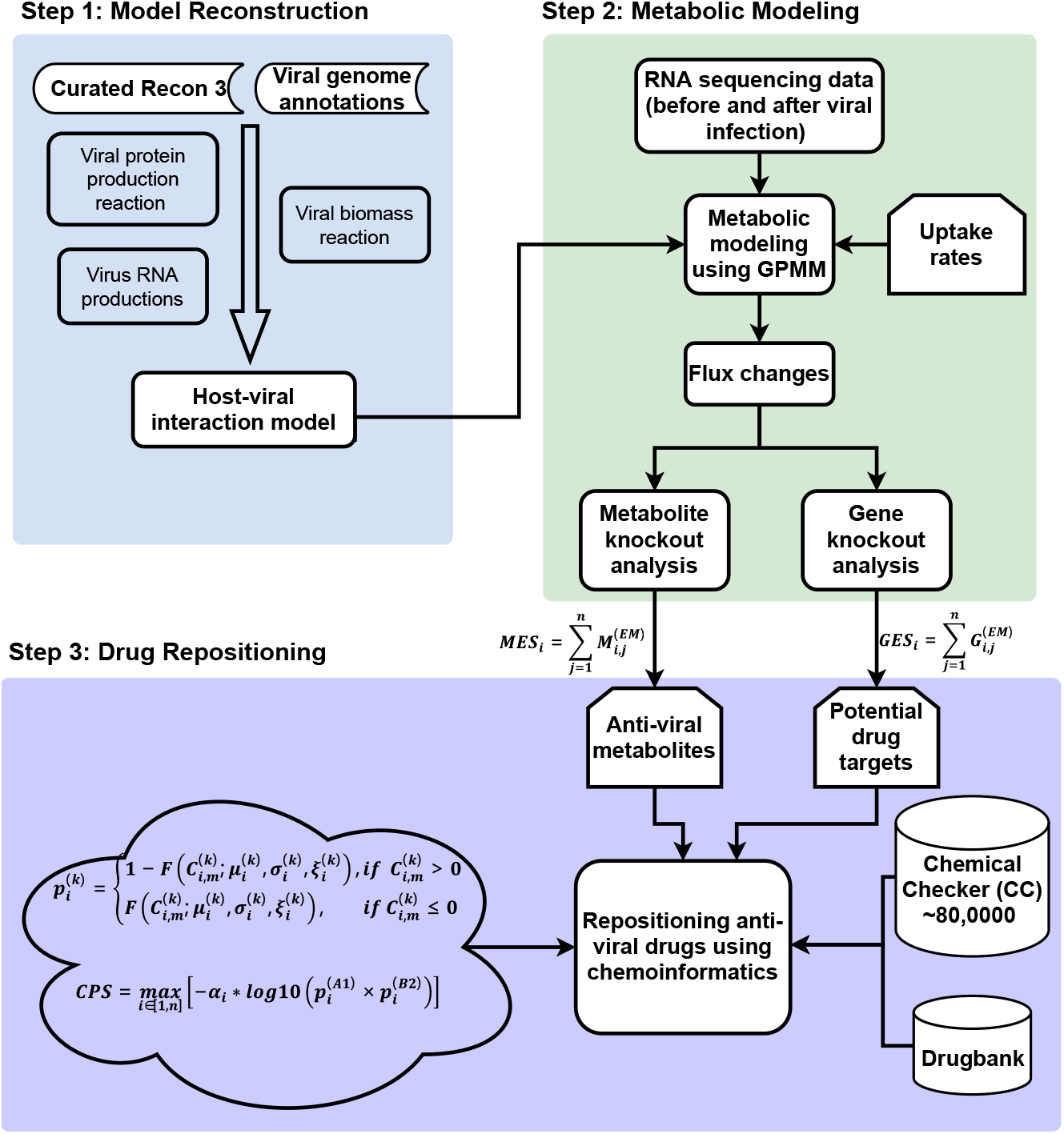
A Flowchart of viral-host Metabolic Modeling (vhMM). Step 1, Model reconstruction of human cells after viral infection. Step 2, Genome-wide precision metabolic modeling to identify metabolic changes after infection and in silicon genome-wide knock-outs to identify genes and metabolites as targets for modulation. Step 3, Drug repurposing by identifying candidate drugs that mimic the molecular characteristics of the gene and metabolite targets.

Second, Genome-wide Precision Metabolic Modeling (GPMM), a method we recently developed to quantitatively integrate enzyme kinetics information from curated knowledge bases and enzyme levels from transcriptome and proteome data into metabolic models[19], was applied to characterize changes in metabolic fluxes due to the viral infection by comparing transcriptome data before and after SARS-CoV-2 infection in three cell lines, A549, Calu3 and NHBE[26].

Third, in silico knockout analysis was systematically performed on all the genes and metabolites involved in the model to examine which genes and metabolites can reverse the altered metabolic fluxes in infected cells, using a computationally efficient algorithm FastMM[27]. Since the virus is critically dependent on redirecting many host metabolic pathways for its reproduction in the infected host cells, different from previous studies that only considered the viral biomass reaction, we considered all the metabolic fluxes changed by the infection to predict potential interventions to restore the compromised host metabolism toward homeostasis. This all-against-all knockout analysis gave rise to effect-size matrices of genes and metabolites on metabolic fluxes from which the top genes and endogenous metabolites were identified as potential targets.

Finally, we adapted Chemical Checker (CC)[23], a database of biological and chemical signatures of ∼800,000 small molecules, and DrugBank[28], an online database containing information on drugs and drug targets, to predict candidate drugs that mimic the molecular characteristics of the gene and metabolite targets. If two compounds have similar metabolic networks or signaling pathways, we define them as having similar effects if they have the same pharmacological effect; otherwise, they are defined as having opposite effects (Details in **Materials and Methods**).

### Evaluation of the host metabolism hijacked by SARS-CoV-2 in infected cells

We analyzed the altered metabolic state of the host cells infected by SARS-CoV-2 using a transcriptome dataset of three viral-infected cell lines (ACE2-induced A549, Calu3, and NHBE)[26]. First, using the viral biomass reaction as the target function, we simulated from the transcriptome data the viral production of SARS-CoV-2 under each of the 13 conditions across the three cell lines. As shown in **Figure 2A**, compared with the measured viral load, the algorithm effectively predicted the viral production (r = 0.75, p = 0.003). Next, we identified individual metabolic fluxes and pathways significantly perturbed in each of the cell lines (false discovery rate < 0.01). Notably, we observed a core set of viral perturbed metabolic pathways changed consistently between the three cell lines after SARS-CoV-2 infection (**Figure 2B**). Strikingly, pathways intimately associated with viral production, such as SARS-CoV-2 protein synthesis, sense RNA synthesis, antisense RNA synthesis, and the overall viral biomass, were all significantly up-regulated, while pathways of oxidative phosphorylation (OP), mitochondria transport, glycolysis/gluconeogenesis, and tyrosine metabolism were down-regulated in the aforementioned three cell lines, presumably as the result of the viral reprioritization of the host metabolism to support the synthesis of viral components. Importantly, these results were corroborated in a most recent study that core mitochondrial genes are down-regulated in nasopharyngeal and autopsy tissues of patients after SARS-Cov-2 infection[29].

**Figure 2:**
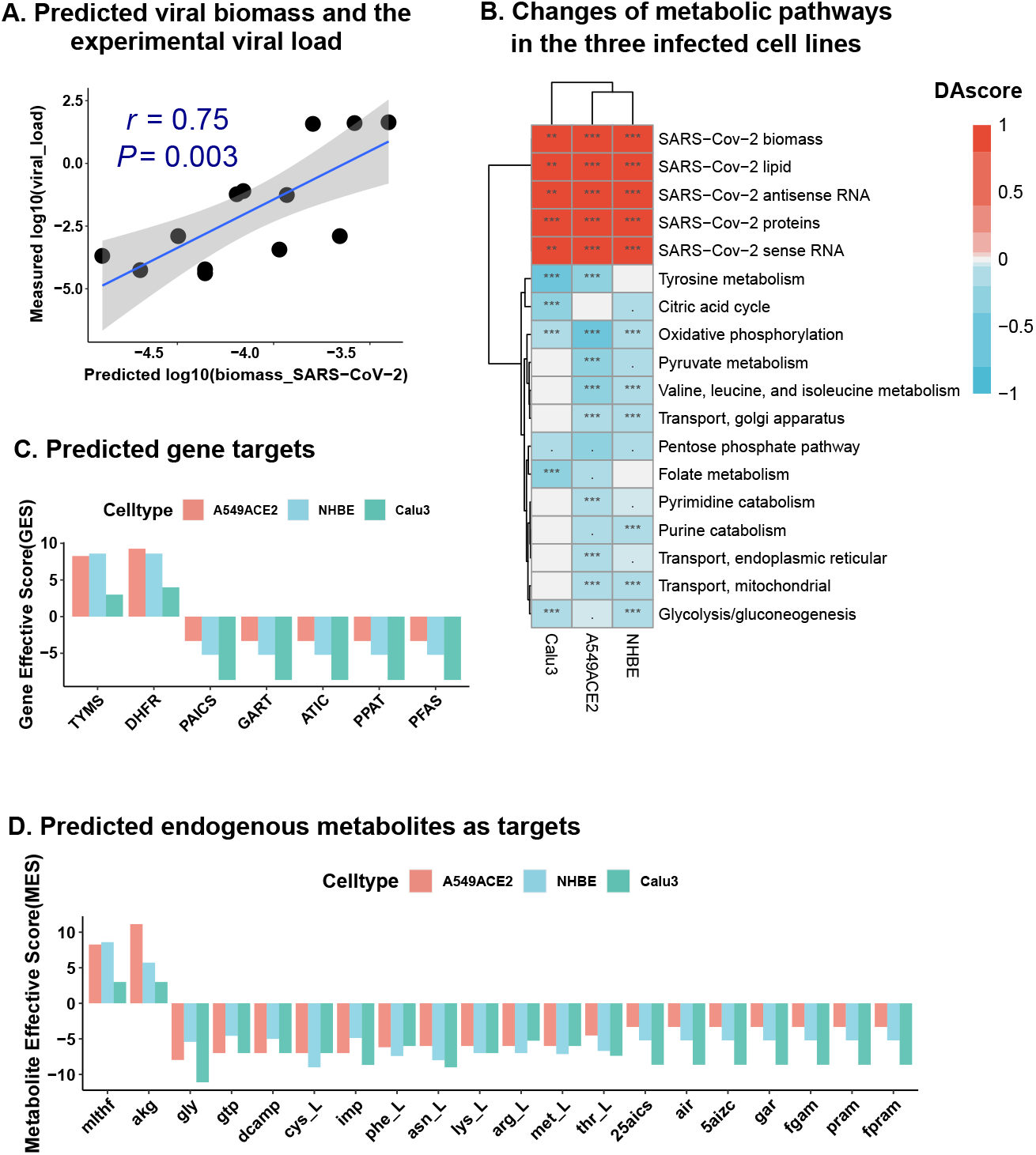
Prediction of gene and endogenous metabolite targets. (A) Correlation between the predicted viral load by vhMM and the experimentally measured viral load (r = 0.75, p = 0.003). (B) Heatmap of changes of metabolic pathways after SARS-COV-2 infection in three cell lines (Calu3, ACE2-induced A549, and NHBE). Note: ***, **, *, and. represent the false discovery rate (FDR) of <0.001, <0.01, < 0.05, and < 0.1, respectively. (C) Gene effective scores of predicted gene targets in the three infected cell lines. (D) Metabolite effective scores of predicted endogenous metabolites as targets in the three infected cell lines.

### In silico gene knockouts toward restoring host metabolism after infection

We performed an all-against-all genome-wide single-gene knockout analysis in the three cell lines and computed changes in metabolic fluxes affected by each gene knockout. These resulted in a matrix of significant gene-flux effects for each cell line that included genes and corresponding metabolic fluxes significantly changed in the cell line (log2 fold changes of the fluxes > 0.5 and p <0.05). To be stringent, we only considered gene knockouts that significantly affecting at least three separate fluxes as potential candidates for targets (Details in **Materials and Methods**). In A549, this matrix comprised 181 genes and 213 fluxes, while 166 genes and 228 fluxes in Calu3 and 171 genes and 154 fluxes in NHBE (**Supplementary Table S1-S3**).

We further defined the genes identified as antagonist targets, where the knocked-outs of these genes inhibited the fluxes up-regulated in the viral infection, and as agonist targets, where the gene knocked-outs further decreased the fluxes down-regulated in the viral infection. We hypothesized that inhibiting the antagonist targets or activating the agonist targets helps restore the infection-altered metabolic fluxes toward homeostasis and thus may reduce viral production. Five antagonist and two agonist targets were identified that were consistent in the three cell lines (A549, Calu3, and NHBE) (**Figure 2C, Supplementary Table S4**). The five antagonist targets were PAICS, GART, ATIC, PPAT, and PFAS, which are all enzymes involved in purine synthesis. Purine synthesis has been widely considered a target for developing antiviral and anticancer drugs[30, 31] and can be potentially targeted against SARS-COV-2 infections. The two agonist targets were TYMS (thymidylate synthetase) and DHFR (dihydrofolate reductase).

We investigated whether these gene targets can directly inhibit the major components of viral production, including viral proteins, RNA, lipids, and the viral biomass reaction. The result showed that knockouts of each of the five antagonists reduced every viral component, including viral proteins, RNA, and lipids, as well as the viral biomass reaction in Calu3 and NHBE cell lines (**Supplementary Table S5)**. For the A549 cell line, knockouts of these antagonist targets inhibited viral proteins and RNA, but not viral lipids and the biomass reaction (**Supplementary Table S5)**. These results indicate that targeting these antagonists could inhibit viral production.

### In silico screening of endogenous metabolites to modulate host cells after infection

We next conducted an all-against-all metabolite knockout analysis to obtain the potential endogenous metabolites to modulate. Similar to the gene knockout analysis, we only considered metabolite knockouts that significantly affected at least three separate fluxes as potential candidates for targets (Details in **Materials and Methods**). 462 metabolites were identified as significantly affecting the metabolic fluxes in A549 infected cell line, while 455 and 400 metabolites were identified as significant in Calu3 and NHBE cell lines, respectively (**Supplementary Table S6-S8**).

These metabolites were also characterized into two categories based on their impact on the metabolic fluxes compared with the corresponding changes by the infection. **Figure 2D** and **Supplementary Table S9** showed the 20 endogenous metabolites identified as consistent in the three cell lines. Two agonist endogenous metabolites included methylenetetrahydrofolate(mlthf), and 2-oxoglutarate (akg), and 18 antagonist metabolites included IMP, GTP, adenylosuccinate (dcamp), seven metabolites related to purine synthesis (25aics, air, 5aizc, gar, fgam, pram, and fpram), and eight amino acids (Gly, Cys, Phe, Asn, Lys, Arg, Met, and Thr). IMP was previously identified as an essential metabolite for viral production, and its synthesis has been targeted in antiviral applications [32].

We further investigated whether targeting these metabolites can also directly inhibit viral production. The results showed that targeting any of the 18 antagonist metabolites can not only inhibit the biosynthesis of all the viral components as well as the viral biomass reaction (**Supplementary Table S10)**.

### Common drug candidates between the original strain and variants of concern (VOCs)

Numerous variants of SARS-CoV-2 have emerged since the pandemic, and so far, at least VOCs have significantly impacted global public health, therapeutics, and vaccines[33].

Therefore, we evaluated whether our results from the study of an original SARS-CoV-2 isolate (IC19) are replicable in independent studies of the different viral isolates (Alpha, Delta, and Omicron) in a cell line (Calu3)[34, 35].

The core set of metabolic pathways identified in IC19 infected cell lines was similarly perturbed by the VOCs (**Figure 3A**), which included the up-regulated SARS-CoV-2 proteins, sense RNA, antisense RNA, and overall SARS-CoV-2 biomass, and the down-regulated oxidative phosphorylation (OP), mitochondria transport, folate metabolism, pyruvate metabolism, and nucleotide interconversion. The overall correlation between Alpha, Delta and Omicron variants and the original IC19, respectively of the changes on these pathways was 0.60, 0.59, and 0.39, which are consistent with the order in which the variants emerged.

**Figure 3:**
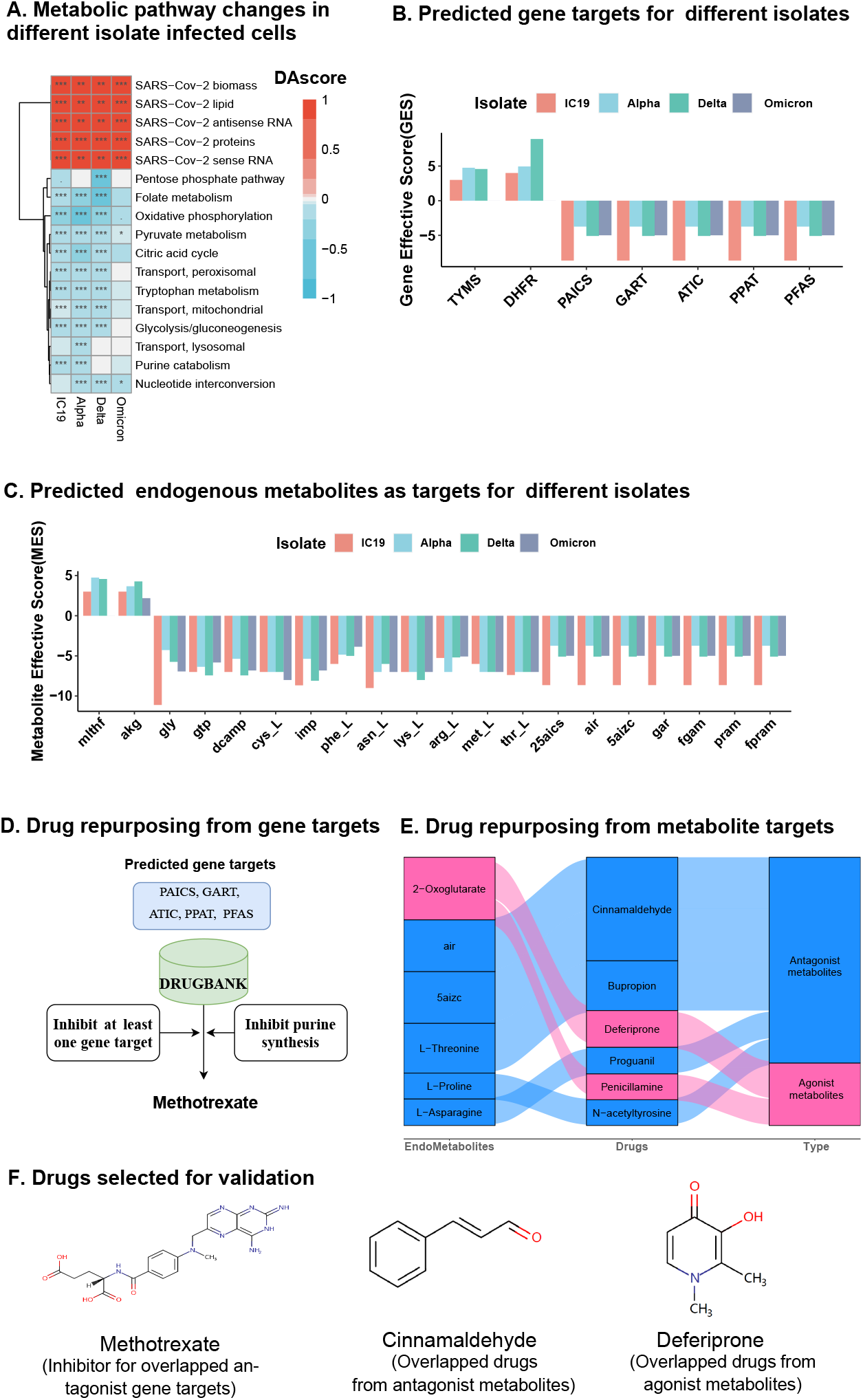
Prediction of repurposing drugs among IC19, Alpha, Delta and Omicron variants. (A) Heatmap of metabolic pathway changes in IC19, Alpha, Delta, and Omicron variant infected cells. Note: ***, **, *, and. represent FDR of <0.001, <0.01, < 0.05, and < 0.1, respectively. (B) Gene effective scores of predicted gene targets in different viral isolates. (C). Metabolite effective scores of predicted endogenous metabolite targets in different viral isolates. (D) Drug repurposing by requiring the inhibition of at least one antagonist gene target and the inhibition of purine synthesis. (E) Drug repurposing based on metabolite targets. Blue and red colors represent antagonist and agonist endogenous metabolites, respectively. The metabolites and predicted drugs are listed in descending order based on the metabolite effective score (MES) and the compound prediction score (CPS), respectively. (F) Drugs selected for in vitro validation. Methotrexate is an FDA-approved drug and is predicted to inhibit the antagonist gene targets and pathway, while Cinnamaldehyde and Deferiprone are the top-ranked drugs from antagonist and agonist endogenous metabolites, respectively.

We next evaluated the metabolic effects of the gene and metabolite candidates for modulation identified from IC19 in the host cells infected by the different viral isolates. For gene targets, except for the agonist gene targets, the five antagonist gene targets (viz., PAICS, GART, ATIC, PPAT, PFAS) were consistent in IC19, Alpha, Delta, and Omicron isolates (**Figure 3B**). For metabolites, 19 of 20 predicted endogenous metabolites (all except mlthf) were consistent from IC19 to omicron isolates (**Figure 3C**).

### Computational drug repurposing for SARS-CoV-2 infection

We first screened the FDA approved drugs based on gene targets. As five consistent gene targets are all purine synthesis enzymes (**Figure 3B**), we screened the FDA-approved drugs that not only inhibit at least one of the predicted target genes, but also have pharmacological effects that inhibit purine synthesis. Methotrexate was identified from the DRUGBANK[28] database that met the above two criteria (**Figure 3D**).

Next, to identify candidates for drug repurposing based on metabolite targets, we overlapped drugs and compounds in DRUGBANK with molecules in Chemical Checker[23], and obtained 2,008 FDA-approved compounds in Chemical Checker. We searched for drugs that have similar network profiles as the endogenous metabolite targets (Details in **Materials and Methods**).

Six of 23 endogenous metabolites can significantly match at least one of the FDA-approved compounds by using network and pharmaceutical similarity (compound prediction score (CPS) > 0 based on network similarity, details in **Materials and Methods**), including 2−Oxoglutarate, 5-Amino-1-(5-Phospho-D-Ribosyl)Imidazole (air), 5-Amino-1-(5-Phospho-D-Ribosyl)Imidazole-4-Carboxylate (5aizc), L−Threonine, L−Proline, and L−Asparagine. At the same time, six FDA-approved drug candidates were identified (CPS>0), including Cinnamaldehyde, Bupropion, Deferiprone, Proguanil, Penicillamine, and N-acetyltyrosine (**Figure 3E**). Notably, 4 of these 6 drugs were previously studied for their antiviral functions (**Supplementary Table S11**).

### *In vitro* experimental validation of the predicted drug candidates

To further test whether the predicted gene target and drug candidates can inhibit viral production, we selected three of the predicted drugs for subsequent validation by *in vitro* cell line experiments. These include methotrexate (an inhibitor of common drug targets), cinnamaldehyde (the top-ranked drug from common antagonist metabolites), and deferiprone (the top-ranked drug from common agonist endogenous metabolites (**Figure 3F**).

We first estimated the drug safety by performing a cytotoxicity assay (Details in **Materials and Methods**). None of the three drugs appeared to have significant cytotoxicity for the drug concentrations up to 100uM (**Figure 4A**). Immunofluorescence also showed that the level of SARS-CoV-2 N protein was significantly reduced at the concentration of 25uM for each of the three drugs(**Figure 4A**).

**Figure 4:**
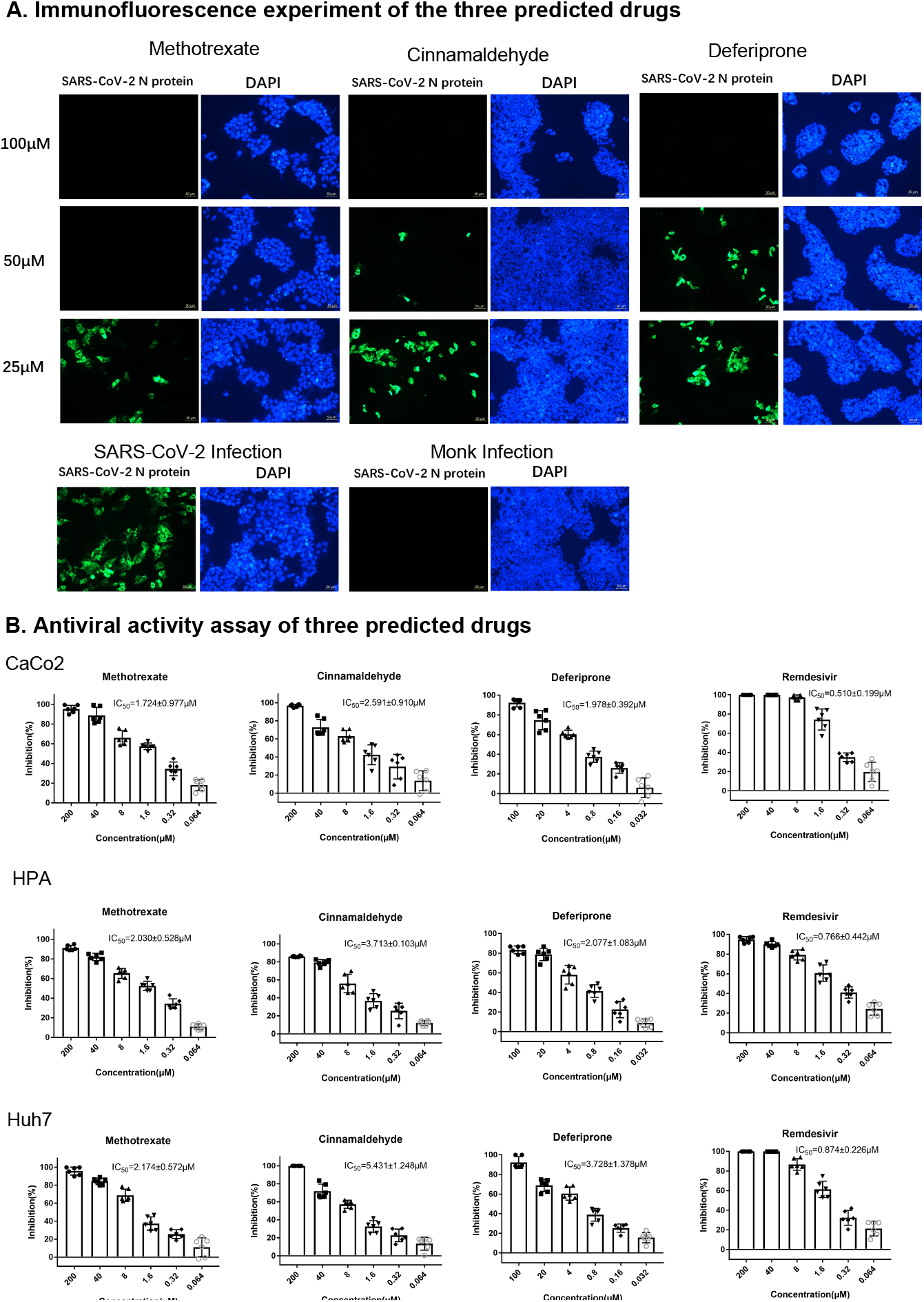
Experimental validation of Methotrexate, Cinnamaldehyde, and Deferiprone. (A) Immunofluorescence experiment of the three predicted drugs. Different drug concentrations (25uM, 50uM, and 100uM) were assayed for each of the drugs. (B) Antiviral activity assay of the three predicted drugs and a positive control (remdesivir). Three lung cell lines (CaCo2, HPA, and Huh7) were used to test the antiviral activity of each drug.

To measure the efficacy in viral inhibition of these three drugs, we performed the antiviral activity assay in three different lung cell types (viz., CaCo2, HPA and Huh7 cell lines) at different dosage range from 0.01uM to 100uM. As shown in **Figure 4B**, the average IC50 of methotrexate among these three cell lines was 2.0 uM (1.72uM for CaCo2, 2.03uM for HPA, and 2.17uM for Huh7). For cinnamaldehyde, the average IC50 was 3.9uM (2.59uM for CaCo2, 3.73uM for HPA, and 5.43uM for Huh7). For deferiprone, the average IC50 was 2.59uM (1.97uM for CaCo2, 2.07uM for HPA, and 3.72 for Huh7). Remdesivir was used as a positive control, which had an average IC50 of 0.83uM as measured (**Figure 4B**).

Together, these results showed that all three tested drugs have a significant anti-viral effect with the IC50 of SARS-CoV-2 ranging from 2.0uM to 3.9uM, clearly supporting that vhMM is effective in computational drug repurposing for anti-viral applications.

### Analysis of the predicted repurposing drugs on real-world patient data

We next attempted to analyze the effect of these three drugs in a real-world setting. Methotrexate had an IC50 of 2.0uM, and the effective dosage was estimated at 40 mg/day (Details in **Materials and Methods)**, which is much higher than the usage of the drug for autoimmune-related diseases (< 20mg/week)[36]. Deferiprone had an IC50 of 2.59uM and an estimated dosage of 55mg/day. However, we could not obtain enough data in ACT Network[37], presumably because deferiprone is an orphan drug for the treatment of iron overload in thalassaemia[38].

Cinnamaldehyde is the main active component in cinnamon (∼3% cinnamaldehyde)[39]. It had an IC50 of 3.9uM and an estimated cinnamon equivalent dosage of 0.59 gram/day. On the other hand, cinnamon supplements typically have a daily dosage of 2-4 grams. We analyzed the incidences of COVID-19 in the ACT Network with respect to cinnamon usage. Since cinnamon is mainly used in supportive care for diabetic patients, we used patients with diabetes as the control group. A retrospective cohort of 410,645 individuals with diabetes from five major hospitals of ACT Network was queried. There was no difference in age distribution between the two groups (**Supplementary Table S12**, Kolmogorov-Smirnov test p-value = 0.166). These included 2,460 patients taking cinnamon, with a rate of recorded SARS-CoV-2 infection of 6.5%, while the rate of infection in the control group of 408,005 patients was 10.1%. Data of the 5 hospitals consistently showed by meta-analysis that patients who took cinnamon had a significant 36% lower rate of SAR-CoV-2 infection (p = 6.1e-8, odds ratio of 0.64 and 95% CI of 0.55∼0.75) (**Figure 5**). These suggest that cinnamaldehyde is a promising candidate for further investigation against COVID-19.

**Figure 5:**
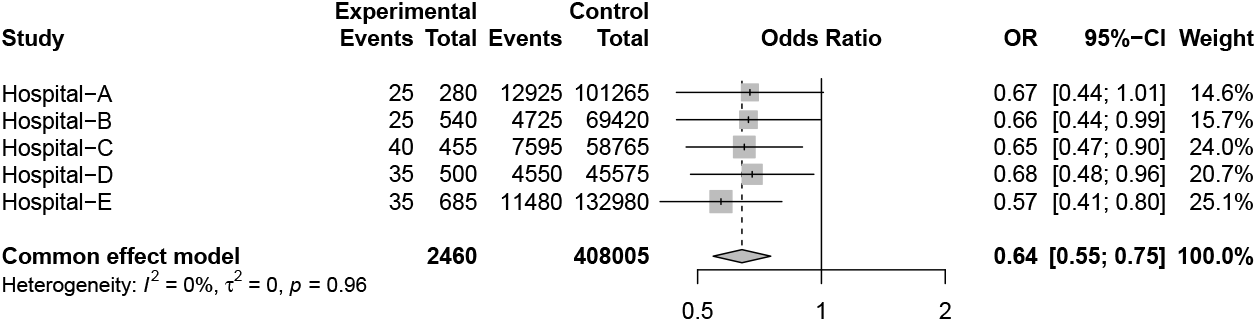
Analysis of a real-world patient dataset on cinnamon. Cinnamaldehyde is the major active component in cinnamon. A retrospective cohort of 410,645 individuals with diabetes from five major hospitals of ACT Network was queried about incidences of COVID-19 with respect to cinnamon usage. These included 2,460 patients taking cinnamon, with a rate of recorded SARS-CoV-2 infection of 6.5%, while the rate of infection in the control group of 408,005 patients was 10.1%. Data from the five hospitals consistently showed that patients who took cinnamon had a significantly lower rate of SAR-CoV-2 infection (p = 6.1e-8, odds ratio of 0.64, and 95% CI of 0.55∼0.75).

## Discussion

In this study, we developed a novel method (vhMM) for drug screening and target identification in viral infectious diseases by integrating metabolic modeling and chemoinformatics. We applied this method to study COVID-19 and captured changes in host metabolism after viral infection. Gene and metabolite targets and six candidate drugs were identified to inhibit both original and mutated SARS-COV-2 production by modulating the host metabolism. We validated the top 3 predicted drugs using *in vitro* experiments and found that all of these candidates can effectively inhibit viral production with IC50 <4uM. In a retrospective cohort study, patients taking cinnamaldehyde showed a significant reduction in the COVID-19 infection rate.

Many viruses, including SARS-CoV-2, depend on the cell metabolism of the infected host for viral replication[40, 41]. vhMM was designed to computationally predict antiviral compounds by modulating the host. In contrast to the commonly used approach of targeting viral proteins[42, 43], vhMM aims to target host metabolism by rescuing or partially rescuing the metabolic dysfunction in host cells after viral infection. We found that the suppression of mitochondria function is one of the major metabolic changes after SARS-CoV-2 infection, which was supported by the recent study of rodent and patient tissues [29]. In addition, the results are consistent in three independent studies from three different laboratories on most pandemic viral variants (alpha, delta and omicron), indicating that vhMM is robust and likely able to tolerate or at least partially tolerate viral mutations. Further, the top candidates were validated by *in vitro* cell line experiments as well as by the analysis of real-world patient data. These results suggest that vhMM is an effective computational method for anti-viral applications.

We performed validation experiments on three predicted candidate drugs, methotrexate, cinnamaldehyde, and deferiprone, and strikingly, all these drugs showed anti-viral activity in multiple lung cell lines. Given that purine synthesis is essential for viral RNA replication and has been approved in other viral diseases[44-46], and a recent study showed that inhibiting the host nucleotide synthesis can effectively block the SARS-COV-2 replication[47], our results further confirmed, at least *in vitro*, that inhibition of purine synthesis by methotrexate can also inhibit the viral production of SARS-CoV-2. Deferiprone is an iron-chelating drug to treat iron overload from blood transfusions in thalassemia[48]. Here we found that deferiprone can also inhibit viral production in a way mimicking 2-oxoglutarate, which may involve modulating cellular redox stress, energy production, or nitrogen metabolism[49, 50].

Cinnamaldehyde, the main active component of Cinnamon, has been previously investigated for its potential beneficial metabolic effects, e.g., increasing glucose uptake and improving insulin sensitivity[51], and cinnamon is sometimes recommended as a supplement for diabetic patients[52]. Here we demonstrated that cinnamaldehyde could inhibit SARS-COV-2 replication by inhibiting intermediate molecules in purine biosynthesis as predicted by vhMM. This is consistent with structure-based prediction of protein targets of cinnamaldehyde in ChEMBL[53]. Retrospective analysis of patient data showed consistent results in patients taking oral cinnamon supplements. Interestingly, the results of a Phase 3 trial (COVID-OUT) recently showed that outpatient treatment with metformin during the initial infection significantly reduced the incidence of long COVID[54], raising the possibility of modulating host metabolism toward homeostasis to prevent the risk of long COVID.

There are still several limitations in our study. First, we did not consider signaling pathways in vhMM, which may result in missing important drug targets and candidates. Second, although the three compounds predicted here have been validated in vitro cell lines, and one of them has been further validated *in vivo* in a large-scale cohort, clinical trials are essential to evaluate their effect in the treatments of COVID-19. Third, the mechanisms of anti-viral activities of these three experimentally supported compounds require further investigation.

## Conclusions

We present a new computational method, vhMM, to evaluate the disturbances of metabolism in virus-infected host cells and to predict repurposing drugs to modulate the host metabolism against viral infections. vhMM was applied to analyze the host metabolism hijacked by SARS-CoV-2 and, importantly, predicted and validated gene and metabolite targets and drug candidates *in vitro* and using real-world data. These results support further drug discovery and repurposing developments to modulate host metabolism in COVID-19 and other viral diseases.

## Methods

### Dataset collections

Datasets from several sources were utilized to develop the computational model for studying the host metabolic response after SARS-CoV-2 infection. These included the genome-wide metabolic model of human cells[55], information on the genomic and proteomic sequences of the SARS-CoV-2 virus[24], RNA-sequencing data of three cell lines (A549, Calu3, and NHBE) infected by the original variant[26], Calu3 cell line infected by the Alpha variant[34] and Calu3 cell line infected by the Delta variant and Omicron variant (BA.5 variant)[35], metabolic uptake dataset[56], chemical information of FDA approved drugs from DrugBank[28], and the harmonized chemical and biological profiles of ∼800,000 small molecules from Chemical Checker[23].

### Genome wide precision metabolic modeling

#### Construction of virus-host metabolic modeling models

First, 28 viral protein production reactions, RNA production reactions, and lipid synthesis reactions were incorporated into the human metabolic model (**Supplementary Table S13**). The viral biomass reaction was then constructed using the structural composition from the experimental data of SARS-CoV-2[25]. The viral biomass reaction is:

2.0 YP_009724393[c] + 1.0 YP_009724397[c] + 0.1 YP_009724392[c] + 0.3 YP_009724390[c] + 0.01 COVID_RNA_BS_pos[c] + 0.01 COVID_RNA_BS_neg[c] + 0.001 COVID_lipids_v[c]

#### Metabolic modeling

We used our recently developed genome-wide precision metabolic modeling method, GPMM, to perform the genome-wide metabolic modeling[19]. Briefly, GPMM integrates the estimated protein abundance from gene expression and enzymatic kinetic parameters into the generic human metabolic model as upper bounds based on Michaelis– Menten kinetics. In addition, the nutrient uptake fluxes of cell lines are derived from the literature[56] and the lower bound of other exchanges was set at zero. The flux variability analysis (FVA) is conducted to construct the tissue-specific models for each cell line using the FastMM toolbox[27].

To obtain the flux change after COVID-19 infection, we performed Monte-Carlo (MCMC) simulations using the Cobra 3.0 toolbox[57]. Since ATP production is essential for all cells, we constrained the low bound of ATP production as 90% of its optimized value, similar to previous studies[58]. Similarly, the lower bound of the fluxes of host biomass and COVID-19 production biomass was set at 90% for the normal and SARS-COV-2 infected cells. Thus, the MCMC sampling provided thousands of feasible fluxes for each reaction but still maintained the feasible ATP production and biomass reaction for the normal and infected cells. The flux changes between normal and infected cells were calculated using the Limma package [59] based on the averaged values of the MCMC results.

#### Identifying metabolic pathway changes

The differential abundance score (DA score) was calculated using the previously published method[60]. For each metabolic pathway (i), the DA score (DA_i_) can be calculated as following:

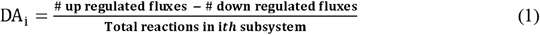

#### All-against-all gene knockout analysis

*In silico* knockout analysis was performed to obtain the effect of each gene knockout on each reaction by using the function of “FastMM_singleGeneKO” in our recently developed FastMM toolbox[27]. We thus obtained an all-against-all gene knockout matrix (G^KO^), where rows and columns represented the genes and reactions, respectively. As the flux of transport was not constrained in GPMM due to the lack of enzymatic parameters[19], we removed the knockout results of transporters.

#### Identification of gene targets

To identify the gene targets for modulation, we first constructed a gene effective matrix(G^(EM)^), which can be calculated as:

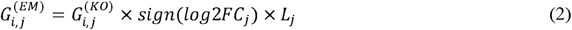

where *i* and *j* represent the *i*th gene and the *j*th gene and the *log*2*FC*_*j*_ and *L*_*j*_ are the log2 fold change and logical of (significance p<0.05 AND log2 fold change >0.5) in the jth flux, respectively.

Then, we calculated the gene effective score (GES) by summing the effective values in G^(EM)^ and calculated as the following:

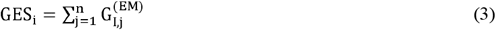

From equation (2), the value of GES >0 or <0 indicates that knocking out this gene can either rescue or enhance the flux changes in the host cells due to infection. To be stringent, we only considered GES > 3 or GES < -3 as potential candidates for antagonist targets or agonist targets, respectively. To obtain robust results, the candidates from each of the three cell types (A549, Calu3 and NHBE) were compared, and only those identified in all the cell lines were designated as gene targets for modulation.

#### All-against-all metabolite knockout analysis

The knockout analysis was conducted using the function of “FastMM_singleMetKO” in the FastMM toolbox to obtain the all-against-all metabolite knockout matrix(M^(KO)^), where row and column represent the gene and reaction, respectively.

#### Identification of endogenous metabolite targets

Similar to the gene targets, we obtained a metabolite effective matrix(M^(EM)^), which was calculated as:

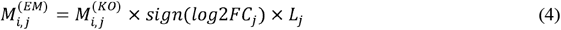

Where 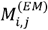 represented the effect score of the ith metabolite on the jth reaction.

Then, we calculated the overall metabolite effective score (MES) by summing the effective values in M^(EM)^:

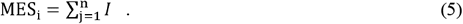

Similar to the gene targets, we only consider the MES > 3 or MES < -3 as the antagonist or agonist candidates, respectively, and the final list of metabolite targets was obtained by also limiting to the candidates consistently found in each of the three cell types.

### Drug repositioning using chemoinformatics

To identify candidate drugs modulating the host cellular metabolism, we reason drugs that have similar network and pharmacological profiles of agonist metabolites or opposite profiles of antagonist metabolites can potentially rescue the COVID-19 metabolic change.

First, we mapped each of the antagonist and agonist metabolites to the Chemical Checker (CC) database and obtained the normalized the fingerprint for 25 different types of molecular features[23]. For each gene target, compounds in CC were also identified that either inhibit or activate the gene, which constitutes the profile of the gene.

Second, we calculated the fingerprint correlations between these metabolites and the ∼800,000 small molecules in CC database to obtain the background correlation distribution. This correlation matrix can be written as 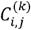, represented the Pearson correlation value of *i*th metabolites and *j*th molecules in kth feature. The value of 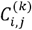 ranges from -1 to 1, where the negative and positive values represent the opposite and similar feature between the two molecules. We used the maximum likelihood method (‘fgev’ function in R ‘evd’ package) to fit the generalized extreme value distribution and estimated the location parameter μ, the scale parameter σ and the shape parameter ξ for each metabolite for each molecular feature. The extreme value distribution of background has been used in our previously developed anti-cancer prediction method [61].

Third, for a given molecule *m*, the one-tailed similarity p-value between m and the screened metabolite *i* 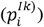 can be calculated as following:

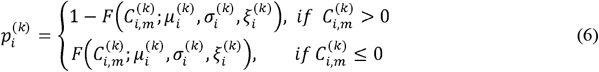

Where 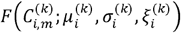 represents the cumulative function of generalized extreme value distribution. 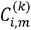 represents the fingerprint correlation between the give molecule *m* and the metabolites *i* in the feature k. 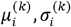 *and* 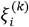 represent the location, scale and the shape parameters of the generalized extreme distribution.

Finally, for each drug compound, we calculate the compound prediction score (CPS) to rank the similarity between the drug compound and the gene and metabolite targets based on three network features: mechanism of action of the compounds (B1 in CC), and pathway features (C2 and C3 in CC), which can be calculated as:

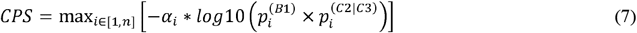

Where *n* is the total number of antagonist and agonist metabolites, *α*_*i*_ is a discriminate factor. When both 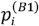 and 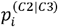 are smaller than p-cutoff (e.g., 0.01), *α*_*i*_ = 1, else *α*_*i*_ = 0.A compound with the CPS >0 was considered a candidate for repurposing.

We retrieved the FDA-approved compounds from DrugBank[28]. After filtering the drugs that were not included in the CC database, we obtained six candidate drugs for anti-COVID-19.

#### *In vitro* experimental validation

The SARS-CoV-2 (NMDCN000HUI) was propagated and titrated on Vero E6 cells (ATCC, no. 1586). Human pulmonary alveolar epithelial cell line (HPAEpiC) was purchased from the ScienCell Research Laboratory (San Diego, CA). Human hepatocarcinoma cell line Huh7 were purchased from Procell Life Science & Technology Co., Ltd (Wuhan, China). Human colorectal adenocarcinoma cell line Caco-2 was purchased from Cell Bank, Chinese Academy of Sciences. All the cells were cultured in high glucose DMEM medium with 4.5 mM L-glutamine (Gibco) contains with 10% FBS (Gibco) and 1% penicillin–streptavidin (Gibco). All work with SARS-CoV-2 was conducted in a BSL-3 facility at the Key Laboratory of Animal Models and Human Disease Mechanisms of the Chinese Academy of Sciences, Kunming Institute of Zoology (Kunming, China).

#### Cytotoxicity assay

The cytotoxicity of compounds on HPAEpiC cells and Huh7 cells was determined by CCK8 assay. Briefly, 8×10^4^ per well HPAEpiC or 4×10^4^ per well Huh7 cells were respectively seeded in 96-well plates and incubated at 37°C, 5% CO2 overnight. When the cells grew to more than 90% confluence, they were incubated with or without series diluted compounds in 96-well cell culture plates and further incubated for 72 hours. 10 μL CCK8 (Beyotime) was added to each well at 37°C, and one hours later, the optical absorbance was measured by ELx800 reader (Bio-Tek) at 450 nm/630 nm. 50% cytotoxicity concentration (CC_50_) was calculated.

#### Immunofluorescence experiment (IFA)

2×10^5^ cells Caco-2 were seeded on glass coverslips pretreated with TC (Solarbio) in 24-well plates to reach 90% confluency and infected with virus at an MOI of 0.02. Meanwhile, the test compounds were added to the wells. After 1 hour, the supernatant was discarded, and the wells were washed 3 times with PBS and replaced with fresh medium containing drug compounds. After 48 hours, cells were fixed in 4% paraformaldehyde and subjected to IFA.

#### Antiviral activity assay

1.6×10^5^ cells of HPAEpiC or 8×10^4^ cells of Huh7 or Caco-2 were respectively seeded in 48-well plates and grown overnight. HPAEpiC cells, Huh7 cells, and Caco-2 cells were infected at an MOI of 1, 0.1and 0.02. At the same time, the test compounds were added to the wells with different concentrations. One hour later, the drug-virus mixture was removed, and cells were washed 3 times to remove the free virus with PBS and replaced with fresh medium containing compounds. 48 hours later, the supernatants were collected, and viral RNA was extracted by using a High pure Viral RNA Kit (Roche), RT-qPCR analysis was performed. Viral RNA was quantified by THUNDERBIRD® Probe One-step qRT-PCR Kit (Toyobo). TaqMan primers for SARS-CoV-2 are 5’-GGGGAA’TTCTCCTGCTAGAAT-3’ and 5’’CAGACA’TTTGCTCTCAAGCTG-3’ with S’RS-CoV-2 probe FAM-TTGCTGCTGCTTGACAGATT-TAMRA-3’. The I’50 values were calculated by using a dose-response model in GraphPad Prism 7.0 software.

#### *In vivo* retrospective cohort study of the ACT network

A rough estimation of the dosage of the repurposed. The drug dose was estimated using the formula by 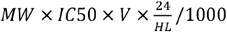, where MW is the molecular weight (unit g/mol), V is the volume of liquid in adults (12L for 60 kg adult), HL is the half-life time (in hours) of the given drug. IC50 is the half maximal inhibitory concentration (in uM). For methotrexate, HL is approximately 6.5 hours[28], and we estimated the dose of 40 mg/day 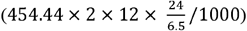. For deferiprone, the HL is 1.9 hours [28], and the estimated dose was 55mg/day 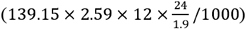. For cinnamaldehyde, the HL is 8.7 hours[62], and the estimated dose is 17.06 mg/day 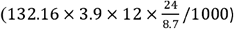. As cinnamaldehyde is the major active component in cinnamon (2.89% cinnamaldehyde)[39], the estimated cinnamon dose is 17.06 mg/day /0.0289 = 0.59 gram/day.

This retrospective cohort study utilized the real-world patient data of the Accrual to Clinical Trials (ACT) network of 35 Clinical and Translational Science Awards (CTSA)-affiliated hospitals[37]. Since cinnamon is mainly used in supportive care for diabetic patients, we queried among patients with a diagnosis of diabetes between 01/01/2020 and 09/09/2022 whether they used cinnamon and whether they had a confirmed case of COVID-19.

The query criteria were the following: i) Diabetes: ICD-9: 250, ICD-10: E08-E13. ii) COVID-19: confirmed cases under CDC case definition, i.e., diagnostic laboratory tests results as active infection test positive. iii) Cinnamon usage: Chromium Picolinate / Cinnamon Bark (1313956), Cinnamon Allergenic Extract (899673), Cinnamon Bark (477053), Cinnamon Preparation (285245).

To obtain robust results, we only included in the analysis data from five hospitals where at least 200 patients took cinnamon. In total, 410,645 individuals with diabetes from these five hospitals were studied: 2,460 patients who took cinnamon were in the case group, and the rest were in the control group. For each hospital, the COVID-19 infection rates were calculated for the case and control groups, and meta-analysis was conducted to estimate the effect of cinnamon usage on COVID-19 infection rate.

## Availability of data and materials

The code and processed data in this manuscript can be available from http://github.com/GonghuaLi/vhMM.

## Acknowledgments

This work was supported by grants from Open Medicine Foundation and the Stupski Foundation (WX), the CAS Project for Young Scientists in Basic Research (YSBR-076), Yunnan Applied Basic Research Project (202101AS070058), High-level Talent Promotion and Training Project of Kunming (Spring City Plan; 2020SCP001), National Key R & D Program of China (2022YFC2303700), Yunnan Key Research and Development Program (202103AC100005) and Yunling Scholar of Yunnan Province.

## Author contributions

GHL, QPK, and WX conceived and designed the study. GHL and WX designed the computational method. GHL developed the computational method, and GHL and FH collected the datasets and performed the analysis. YTZ designed the validation experiments, and RHL and XYL conducted the validation experiments. PL, GHL and WX performed the analysis of real-world data. GHL, FH, CJC, and WX performed the statistical analysis. GHL, FH, RHL, PL, and WX drafted the manuscript. WX, GHL, FH, RHL, PL, WHX, JFH, QPK, and YTZ revised the manuscript. All authors read and approved the final manuscript.

## Ethics declarations

### Ethics approval and consent to participate

Not applicable.

### Consent for publication

Not applicable.

